# Membrane Dynamics-Mediated Heat Retention Dictates Economical Intracellular Energy Flow

**DOI:** 10.64898/2026.04.25.720769

**Authors:** Akira Murakami, Tasuku Sato, Haruya Suzuki, Takashi Funatsu, Kohki Okabe

**Author notes:** Correspondence should be addressed to: K.O.

## Abstract

Intracellular thermometry has revealed temperature gradients within living cells, indicating the presence of localized energy flow. Moreover, increasing evidence suggests that cells can autonomously regulate heat production and use this heat to support cellular functions. Although heat is generally assumed to dissipate rapidly, recent studies suggest that slow, non-diffusive heat dissipation can generate high local temperatures, implying that heat may be retained within cells. However, such intracellular heat retention at the single-cell level has not been experimentally examined. Here, we measured the dissipation of metabolically generated heat using calorimetry and evaluated cellular heat balance. We further separated the dissipated heat into heat transfer and heat production components using an intracellular temperature imaging-based method. Through this approach, we found that perturbation of the cell membrane accelerated intracellular heat transfer rates and increased heat dissipation. These findings provide the first experimental evidence of intracellular heat retention, showing that heat generated by catabolic metabolism is not fully dissipated but is partly retained within cells by the cell membrane, which separates the intracellular space from the external environment. Furthermore, we identified the mechanism underlying this heat retention, demonstrating that membrane lipid dynamics are a key molecular determinant and that the motions of membrane lipids form an energetic barrier to heat dissipation. Together, our findings redefine metabolic heat, long regarded as mere waste, as a functional resource that supports efficient energy use, and they establish membrane-controlled heat retention as a fundamental principle of bioenergetic optimization in living systems.

## Introduction

Animals acquire energy by catabolizing nutrients from their external environment. The energy gained is harnessed to drive physiological functions and is ultimately dissipated as heat. Determining how the bioenergetics of metabolism govern the life cycle is a central quest in biology^1–3^. Chronic disruption of the energy balance (how much energy is gained and dissipated) by obesity or aging results in severe diseases that have long plagued humanity^4^. In particular, for intractable diseases, such as cancer, the collapse of bioenergetics is being extensively researched as a critical determinant of pathogenesis^5^. Thus, elucidating the mechanisms underlying energy balance is of biological, medical, and societal significance and has been a central focus of scientific research.

Traditionally, energy balance has been studied at the whole-organism level. However, a deeper understanding requires investigating it at the cellular level—the fundamental unit of life. The energy derived from cellular catabolism is partitioned into adenosine triphosphate (ATP) and thermal energy. Although the dynamics and functions of intracellular ATP have been extensively characterized across diverse biological contexts, technical limitations have hindered direct measurement of heat flow at the cellular level. Consequently, the full extent of thermodynamic energy flows that constitute the essence of energy balance remains elusive. In light of this gap, recent advances in intracellular thermometry have begun to shed light on heat dynamics in living cells. Notably, we developed a molecular nanothermometer based on a fluorescent polymer that revealed spatiotemporal variations in intracellular temperature of several degrees Celsius^6^. Furthermore, nanothermometers operating on distinct principles have detected temperature variations at the organelle level, including in the mitochondria^7,8^, endoplasmic reticulum (ER)^7,9^, lysosomes^10,11^, and nucleus^12,13^. These variations have been observed not only in thermogenic cells, such as brown adipocytes, but also across a wide range of cell types. Together, these findings indicate the existence of intracellular temperature gradients and imply the presence of heat flow within cells. We further discovered that cells exploit endogenously generated heat as a means of signaling, which we refer to as thermal signaling. For instance, in a brain slice model of edema, activation of the glutamate receptor during neuronal activity leads to an increase in tissue temperature. This, in turn, activates the temperature-sensitive ion channel TRPV4, triggering brain swelling^14^. More recently, we demonstrated that during nerve growth factor (NGF)-induced neuronal differentiation, transcription-dependent increases in intracellular temperature drive neurite outgrowth^15^. Although heat has long been regarded as a metabolic byproduct, these findings indicate that intracellular processes are not driven solely by ATP but also by intracellularly produced heat (i.e., emerging thermal signals). This paradigm shift suggests that the cellular energy balance is not merely a metabolic indicator but also a phenomenon possessing fundamental physiological significance that shapes cellular programs.

However, the fate of the heat generated within the cells remains unclear. If the heat within the cells is dissipated only by conduction, as in homogeneous aqueous solutions, the heat generated metabolically is assumed to be dissipated rapidly^16^. This conceptual tension has given rise to the emerging field of intracellular heat transfer, which combines artificial heating with nanothermometry^17–20^. Using infrared laser–based localized heating, we recently developed a real-time imaging technique that visualizes changes in temperature distribution within living mammalian cells. This technique reveals an unconventionally slow, nondiffusive heat transfer during continuous heating on the scale of seconds^20^. Because spontaneous, metabolism-driven heat generation occurs continuously, these observations suggest that the heat generated within the cells is locally retained rather than rapidly dissipated. These phenomena appear to provide the first plausible explanation for the large intracellular temperature gradients and associated thermal signaling. However, the mechanism underlying non-diffusive heat transfer remains unclear, largely because the fate of metabolically generated heat in single cells has never been experimentally tracked from its generation to its eventual dissipation, hindering a complete depiction of cellular energy balance and flow, as well as potential applications, such as the manipulation of thermal signaling.

Calorimetry is a quantitative technique that detects exothermic and endothermic reactions as heat flows under adiabatic conditions. It has long been used to analyze thermodynamic reactions, including state changes of biomolecules such as phase transitions and thermal denaturation. Calorimetry has also been adopted for biological research, such as the measurement of heat flow in living cells. For example, isothermal titration calorimetry (ITC) and isothermal microcalorimetry (IMC) quantify changes in heat flow at a constant temperature^21^. These techniques have been used to analyze protein–protein and protein–ligand interactions, as well as cellular metabolic activity. In contrast, differential scanning calorimetry (DSC) measures temperature-dependent thermodynamic reactions and has been widely used to characterize the thermal properties of biomolecules^22^. Previous DSC studies on living cells have mainly focused on detecting heat denaturation or evaluating cell viability under thermal stress^23^. Several studies have also observed catabolism-dependent heat production within physiological temperature ranges^24,25^. Thus, DSC is well-suited to our aim of comprehensively capturing and characterizing the total heat associated with chemical reactions occurring across a broad temperature range in living cells, including those arising from metabolism, to understand the cellular energy balance.

The cell membrane shapes the cell and embodies its structural and functional identity of life. Lipids are extraordinarily abundant within cells, a fact that is often overlooked, because they constitute not only the plasma membrane, which separates the cell from its surroundings, but also the membranes that divide the intracellular space into compartments. The physicochemical properties of the cell membrane, such as fluidity, are highly dependent on temperature and can be precisely adjusted by molecular composition. Many organisms adapt to changes in environmental temperature by altering the composition of their cell membranes^26^. For example, an increase in unsaturated fatty acids in the phospholipid acyl chains enhances membrane fluidity. This is a long-recognized strategy for coping with low-temperature environments. Using single-cell thermometry, we uncovered a previously unknown role of lipid remodeling^27^. During cold exposure, desaturation of phospholipid acyl chains promotes heat production, suggesting that it helps maintain intracellular temperature. These findings reveal a connection between cell membranes/lipids and environmental (macroscale) and intracellular (nanoscale) temperatures. This renders cell membranes and lipids plausible candidates for determining intracellular heat transfer and shaping cellular energy balance.

In this study, we employed DSC to quantitatively analyze cellular energy balance, with a particular focus on heat metabolically generated within cells. We also examined the effect of cell membranes on cellular energy balance. We integrated the cellular thermodynamic insights obtained from DSC with the spatially resolved information provided by our nanothermometry-based quantitative local heating approach. These versatile analyses enabled us to disentangle the heat-retention capacity from cellular heat-production activity and demonstrated that membrane dynamics govern the cellular heat balance by preventing the rapid dissipation of metabolically generated heat. We further revealed the molecular mechanism underlying this process. These findings provide the first experimental evidence of an intrinsic system that enables cells to harness locally generated heat.

## Results

### Microcalorimetry enables quantification of cellular exothermic heat flow from intracellularly generated heat

To understand the cellular heat balance, we employed highly sensitive microcalorimetry based on DSC and quantified the heat flow of the cells. Our DSC instrument contained a sample chamber filled with living cells or cell extracts and a reference chamber filled with the corresponding control solutions, such as medium or buffer. The two chambers were placed in an adiabatic system (Fig. 1a). As the system temperature increased at a constant rate, the temperatures of both chambers were continuously monitored. When an exothermic or endothermic reaction occurred in the sample chamber, a temperature difference (Δ*T*) developed between the two chambers (Fig. 1b). Dividing this Δ*T* by an instrument-dependent calibration constant (*R*) yielded the heat flow of the sample, which was, in principle, proportional to the scan rate (Fig. 1c). When living HeLa cells suspended in culture medium were scanned by DSC from 5°C at a constant heating rate, an exothermic signal increased progressively as the temperature increases to a maximum, around 45°C. Above 45°C, the exothermic signal gradually diminished and transitioned to an endothermic signal that peaked around 65°C (Fig. S1a, 1st scan). Rescanning the HeLa cells that had been thermally denatured during the first scan produced a largely flat heat flow trace with only a slight trend. The third scan yielded a trace identical to that of the second scan (Fig. S1a, 2nd and 3rd scans). Based on these observations, we subtracted the second scan trace from the first to remove the baseline and background contributions (Fig. S1b, 1st–2nd). The validity of this procedure was confirmed by subtracting the third scan trace from the second, resulting in an almost featureless trace (Fig. S1b, 2nd–3rd).

**Figure 1.**
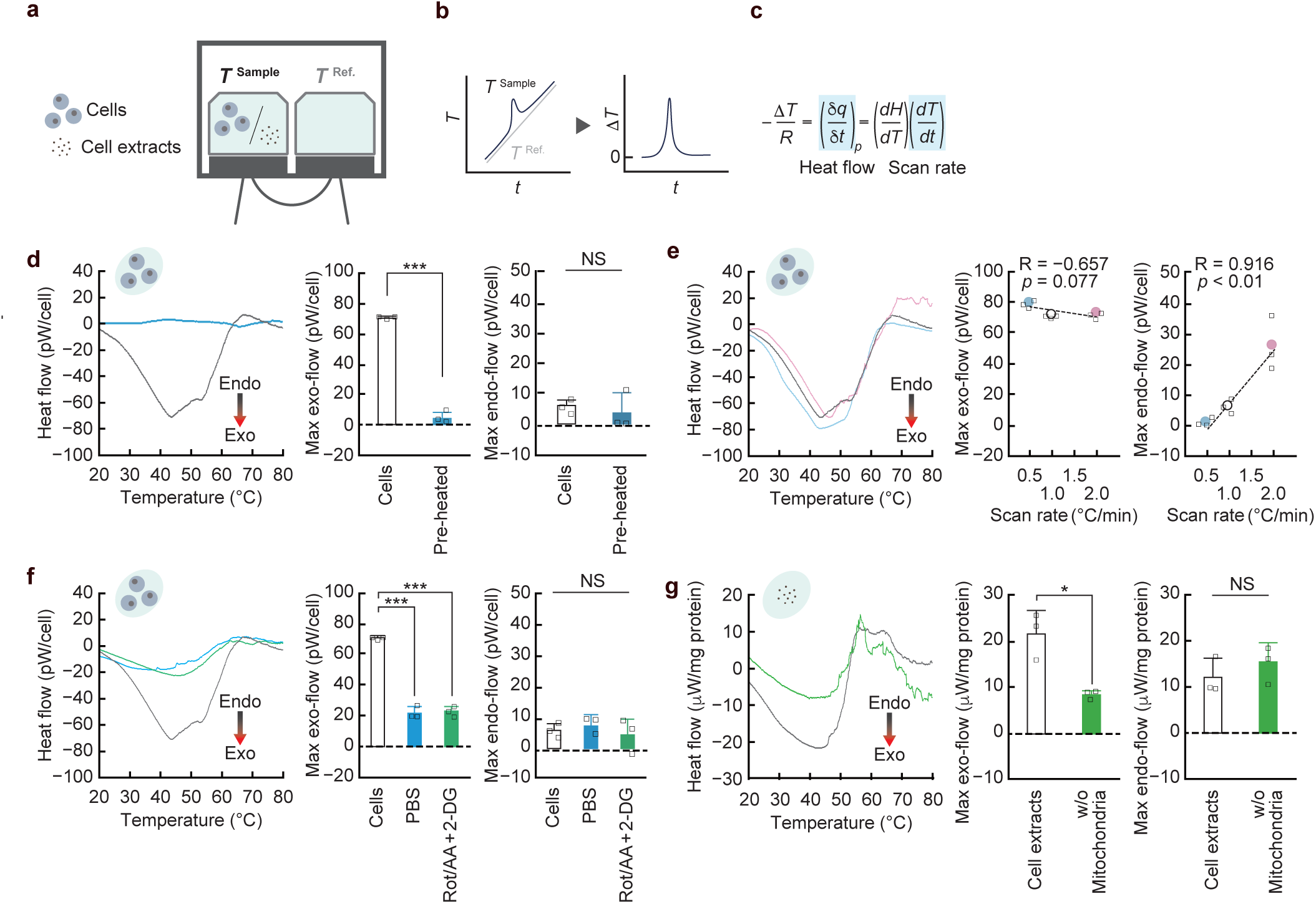
Differential scanning calorimetry (DSC)-based quantification of cellular heat flow. **a.** Schematic illustration of the DSC system. **b.** Time (t)-dependent temperature (*T*) difference (Δ*T*) generated between the sample and reference chambers when an exothermic reaction occurs in the sample chamber. **c.** Schematic explanation of the relationship between heat flow calculated from Δ*T* and the DSC scan rate. **d.** Heat flow in living HeLa cells. The DSC curve was compared with that obtained from preheated cells (*n* = 3). **e.** Effect of scan rate on heat flow measured in living HeLa cells (*n* = 3). The dotted lines indicate linear fitting. R indicates the correlation coefficient; p values were used to assess statistical significance, with *P* < 0.05 considered significant. **f.** Effects of catabolic metabolism inhibition on heat flow in living HeLa cells. Rotenone and antimycin A (Rot/AA) are inhibitors of mitochondrial respiratory complexes I and III, respectively; 2-deoxy-D-glucose (2-DG) is a glycolysis inhibitor (*n* = 3). **g.** Heat flow in HeLa cell extracts. The DSC curve was compared with that of cell extracts without the mitochondrial fraction (*n* = 3). DSC measurements were typically performed at a scan rate of 1.0 °C min^−1^. In the DSC thermograms, mean traces are shown. Negative values on the y-axis represent exothermic heat flow, whereas positive values indicate endothermic heat flow. For bar graphs, the maximum exothermic and endothermic signals in each DSC thermogram were extracted and defined as Max exo-flow and Max endo-flow, respectively. Data are shown as means + standard deviation (s.d.). In **e** and **f**, the DSC data are compared with those obtained from living HeLa cells in d. \**P* < 0.05, \*\*\**P* < 0.001, NS indicates not significant.

First, we characterized the heat flow detected in live HeLa cells, which was almost eliminated by preheating (Fig. 1d). The maximum exothermic flow within the physiological temperature range (35–45°C) was 71.1 pW, which is comparable to previous reports^28,29^. The endothermic peak around 65°C exhibited a strong positive correlation with the scan rate (Fig. 1e). The dependence of the endothermic flow on the scan rate closely resembled that observed for the thermal denaturation of bovine serum albumin (BSA) when measured using our system. This finding is consistent with the aforementioned DSC principles (Figs. 1c and S2). These observations suggest that the endothermic signal originates from the thermal denaturation of abundant intracellular biopolymers, which is consistent with the findings of a previous study^30^. However, the exothermic peak did not exhibit any correlation with the scan rate (Fig. 1e). This suggests that the exothermic signal detected in HeLa cells does not arise from transient temperature-dependent state changes, such as protein denaturation.

We next hypothesized that the scan-rate-independent exothermic signal originated from constant exothermic reactions and examined catabolic metabolism as a potential source of these reactions. The exothermic signal of live HeLa cells decreased significantly when the cells were suspended in phosphate-buffered saline (PBS), a nutrient-deprived medium (Fig. 1f). Glycolysis and the subsequent electron transfer through the mitochondrial respiratory chain play central roles in catabolic metabolism. Consistently, treatment with 2-deoxy-D-glucose (a glycolysis inhibitor) and rotenone/antimycin A (inhibitors of the mitochondrial respiratory chain) reduced the exothermic signal to a level comparable to that observed in PBS, even in nutrient-rich normal culture media (Fig. 1f). Cell extracts prepared from HeLa cells exhibited exothermic flow, which was significantly reduced by the biochemical removal of the mitochondrial fraction (Fig. 1g). These findings indicate that living cells exhibit continuous heat flow from inside to outside the cell at physiological temperatures and that this heat flow depends on metabolic activity, particularly catabolic reactions.

After quantifying the exothermic (i.e., dissipated) heat flow from living cells, we evaluated the quantitative capability of our DSC system by estimating the amount of heat generated through catabolic metabolism and comparing this value with the heat flow measured by DSC. We measured changes in cell number, glucose consumption in the culture medium, and lactate release into the medium (Fig. S3a-c). Over the 24 h culture period, glucose levels in the medium significantly decreased, whereas lactate levels increased significantly (Figs. S3b and 2c). As detailed in the Supplementary Note, using glucose consumption as a readout of total metabolic flux (aerobic and anaerobic) and lactate production as an indicator of anaerobic metabolism, we calculated that HeLa cells generated 89.1 ± 3.6 pW of heat through glucose catabolism. The calculated heat generated (89.1 pW) was on the same order of magnitude as, although somewhat higher than, the exothermic heat flow detected by DSC at 37°C during the scan (49.3 ± 0.7 pW) and the maximum exothermic heat flow (71.1 ± 1.0 pW). These results support the quantitative accuracy of our DSC system. Thus, we developed a DSC-based system that can quantitatively measure the exothermic heat flow constitutively and catabolically generated by living cells.

### The integrity of the cell membrane suppresses heat flow from living cells

We focused on the cell membrane and membrane lipids as potential factors that constrain heat transfer within the cells and the eventual dissipation of heat to the extracellular environment. Cholesterol is a major component of the mammalian cell membrane, particularly the plasma membrane, which separates the inside of the cell from the outside^31^. Thus, altering cellular cholesterol levels can easily and efficiently disrupt the physicochemical properties of cell membranes. In this study, we examined the effect of cholesterol depletion on exothermic heat flow in HeLa cells. Methyl-β-cyclodextrin (MβCD), a cyclic compound, extracts cholesterol from the plasma membrane through inclusion complex formation. Initially, we found that MβCD treatment reduced the fluorescence signal of filipin, a cholesterol indicator, particularly at the plasma membrane. This indicated a decrease in membrane cholesterol content (Fig. 2a). DSC analysis revealed that MβCD-treated cells exhibited increased maximal exothermic heat flow within the physiological temperature range. This effect was suppressed by preloading MβCD with cholesterol, as preloading diminishes MβCD’s cholesterol-extracting activity^32^ (Fig. 2b). In contrast, cell extracts prepared after the MβCD treatment, which lack an intact plasma membrane structure, exhibited DSC profiles similar to those of control extracts (e.g., the same level of exothermic heat flow; Fig. 2c). As the exothermic heat flow detected within the physiological temperature range originates from constitutively and endogenously generated heat in living cells (Fig. 1), these results indicate that cholesterol depletion increases flow of catabolically generated heat from the interior of living cells to the external environment. Under steady-state conditions, the heat dissipation measured by DSC was significantly lower than that estimated from catabolic metabolism (Fig. S3d); these observations raised the possibility that the increased heat flow reflects the release of heat that had previously been retained within the cells in a manner dependent on the integrity of the plasma membrane.

**Figure 2.**
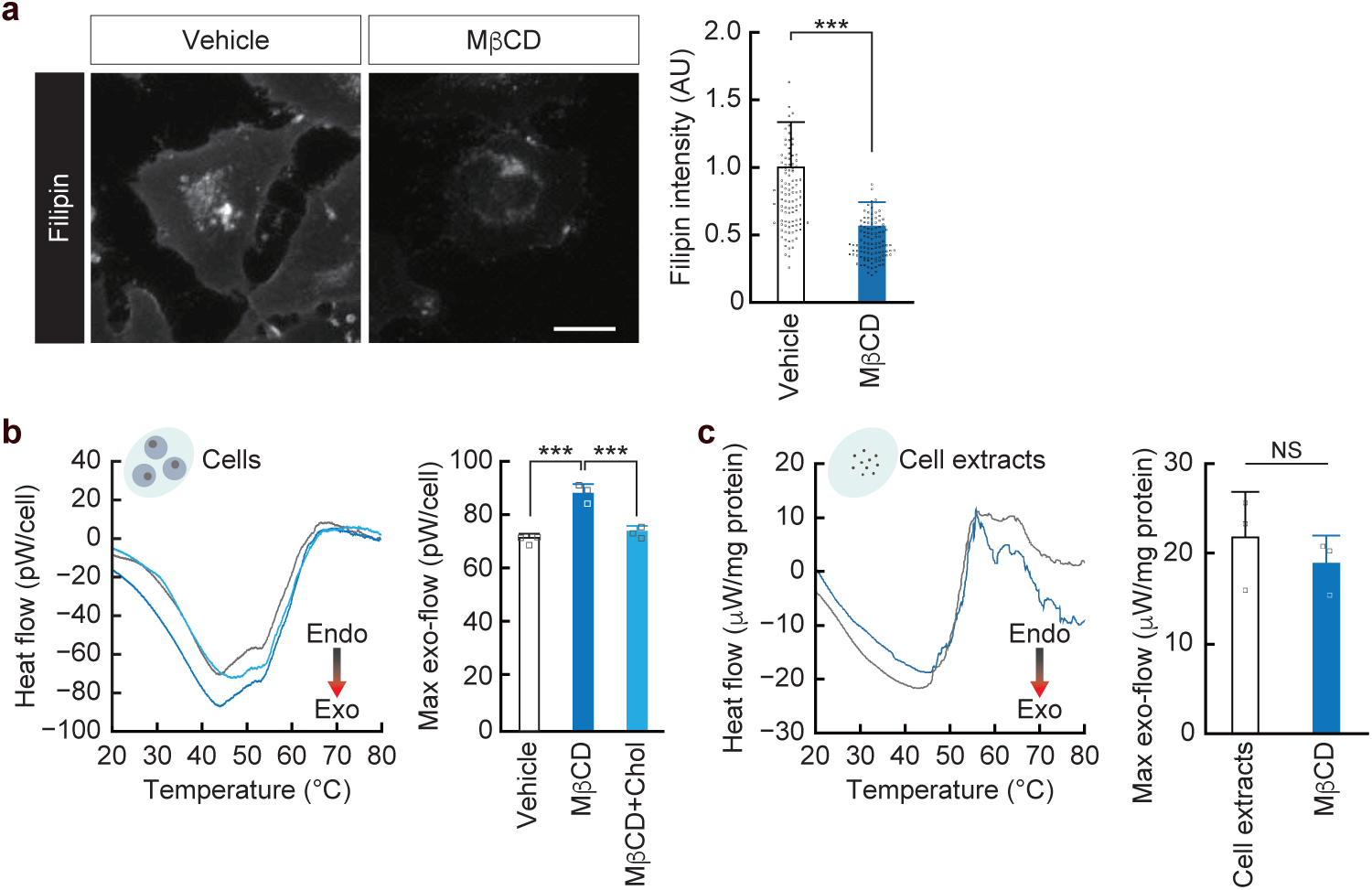
Exothermic heat flow from cells with perturbed plasma membranes. **a.** Effect of MβCD treatment (10 mM for 30 min) on cellular cholesterol levels. Cholesterol was visualized using filipin. Data are shown as means + standard deviation (s.d.) (n = 103 [Vehicle] and 109 [MβCD] cells). Scale bar: 20 μm. **b.** Effect of MβCD treatment on the exothermic heat flow of living HeLa cells. MβCD + Chol indicates treatment with cholesterol-preloaded MβCD (*n* = 3). **c.** Heat flow in cell extracts prepared from HeLa cells with or without MβCD treatment (*n* = 3). In the DSC thermograms, mean traces are shown. Negative values on the y-axis represent exothermic heat flow, whereas positive values indicate endothermic heat flow. For bar graphs, the maximum exothermic signal in each DSC thermogram was extracted and defined as Max exo-flow. Data are shown as means + standard deviation (s.d.). Vehicle indicates treatment with PBS. In **c**, the dataset is compared with that obtained using cell extracts from intact HeLa cells (Fig. 1g). \*\*\**P* < 0.001, NS indicates not significant.

### Quantitative single-cell thermal analysis revealed that the cell membrane restricts heat dissipation from cells

The observed increase in exothermic heat flow in cholesterol-depleted cells may reflect increased cellular heat production. Therefore, we assessed the intracellular heat production at the subcellular level by examining organelle functions that could contribute to heat generation via catabolism. Mitochondria play an essential role in aerobic glucose metabolism via electron transport. The mitochondrial membrane potential, an indicator of electron transport activity, did not change in cholesterol-depleted cells (Fig. S4a). Consistently, the oxygen consumption rate, another readout of respiratory chain activity, also showed no significant difference between MβCD-treated and control cells (Fig. S4b). Subsequently, we evaluated the activity of sarco/endoplasmic reticulum Ca²⁺-ATPase (SERCA), which transports Ca²⁺ from the cytosol into the ER using energy derived from ATP hydrolysis. A portion of the ATP hydrolysis-derived energy is known to be uncoupled from Ca²⁺ transport and instead dissipated as heat^33^. Under Ca²⁺-free extracellular conditions, inhibition of SERCA by thapsigargin (Tg) induces the release of ER-stored Ca²⁺ into the cytosol, thereby increasing cytosolic Ca²⁺ levels; the amplitude of this increase reflects SERCA activity^34^. The Tg-induced increase in cytosolic Ca²⁺ was comparable between MβCD-treated and control cells (Fig. S4c). These results indicate that cholesterol depletion does not alter intracellular catabolic activity. Indeed, intracellular ATP levels did not differ significantly between MβCD-treated and control cells (Fig. S4d).

We then focused on the cellular capacity to retain the generated heat and observed heat dynamics at the subcellular level. For this purpose, we used our previously developed fluorescent polymeric thermometer (FPT)^6^, a temperature-responsive polymer containing hydrophilic side chains and a fluorophore moiety (Fig. 3a). The fluorophore exhibits weak emission when hydrated. However, temperature-induced dehydration, which is associated with conformational changes in the polymer backbone, results in enhanced fluorescence. Fluorescence lifetime imaging microscopy (FLIM) revealed that the fluorescence lifetime of FPT in live HeLa cells increased with increasing temperature of the culture medium (Fig. 3b). These results confirm that intracellular temperature variations can be quantitatively estimated from the corresponding changes in fluorescence lifetime, as previously reported^6,15^.

**Figure 3.**
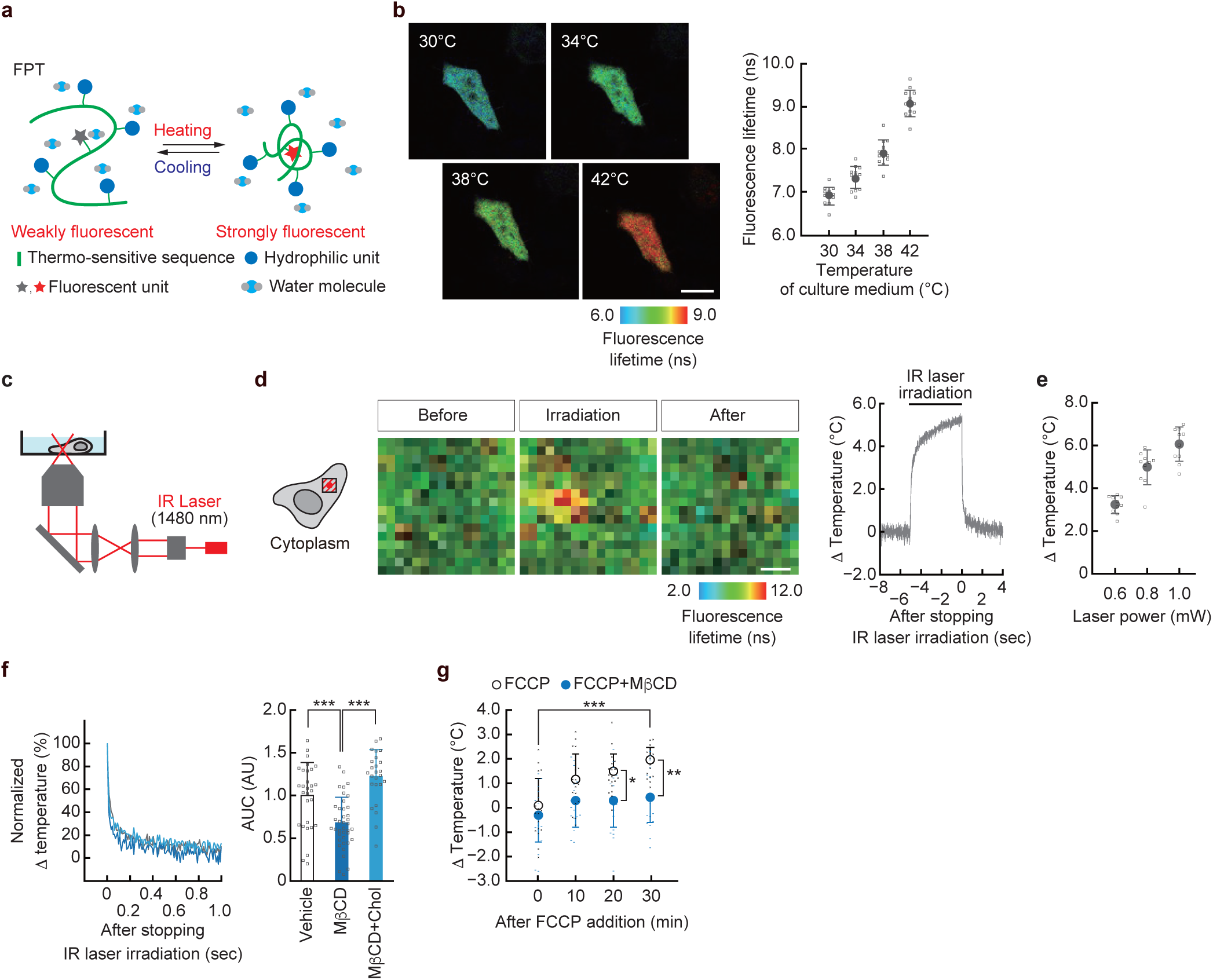
Quantitative single-cell thermal analyses of cells with perturbed plasma membranes. **a.** Schematic illustration of FPT. **b.** Temperature-dependent elongation of the FPT fluorescence lifetime in living HeLa cells. Scale bar: 20 μm. Data are shown as means + standard deviation (s.d.) (n = 13 cells). **c.** Schematic illustration of single-cell temperature manipulation using IR laser irradiation. **d.** Tracking of the average temperature within a square area measuring 10 μm on each side and centered on the heating point during IR laser irradiation (0.8 mW for 5 s; *n* = 29 cells). ΔTemperature represents the temperature change relative to the preheating baseline. Scale bar: 2 μm. **e.** Relationship between IR laser power and the magnitude of ΔTemperature induced in the cytoplasm by 5 s of IR laser irradiation. Data are shown as means + s.d. (*n* = 10 cells). **f.** Effect of MβCD treatment on temperature relaxation after stopping heating (0.8 mW for 5 s). Traces were normalized to the temperature at the time point when irradiation was stopped. Data are shown as means (left). The relaxation rate, an index of intracellular heat transfer, was quantified as the area under the relaxation curve (AUC). Data are shown as means + s.d. (right) (*n* = 29 [Vehicle], 39 [MβCD], and 25 [MβCD + Chol] cells). Vehicle indicates treatment with PBS. **g.** Effect of MβCD treatment on the time-dependent increase in intracellular temperature induced by activation of endogenous cellular heat production. Carbonyl cyanide-p-trifluoromethoxyphenylhydrazone (FCCP), an uncoupler, was used at 5 μM. ΔTemperature represents the change in temperature relative to the value before FCCP addition. Data are shown as means + s.d. (*n* = 19 [FCCP] and 18 [FCCP + MβCD] cells). \*\*\**P* < 0.001.

To quantitatively investigate the effect of plasma membrane perturbation on heat dissipation from cells, we directly observed the intracellular heat transfer dynamics using our previously established approach^20^. In this method, localized cytoplasmic heating was induced by irradiating the cells with an infrared (IR) laser (1480 nm) through an objective lens (Fig. 3c). Intracellular thermometry with FPT confirmed that transient IR laser irradiation induces reversible temperature changes within micrometer-scale regions of the cytoplasm (Fig. 3d) and that the magnitude of the temperature increase depends on the power of the laser and can be quantitatively controlled (Fig. 3e). Using this system, we quantified the rate of intracellular heat transfer by analyzing relaxation kinetics following laser-induced heating. We used the area under the curve (AUC) of relaxation as an index of the relaxation rate; smaller AUC values indicate faster relaxation (Fig. 3f, left). In cholesterol-depleted cells, heat relaxation was significantly faster than that in control cells, reducing the AUC by 32%. Conversely, preloading MβCD with cholesterol abolished this effect (Fig. 3f). These results demonstrate that cholesterol depletion accelerates intracellular heat transfer. As the relaxation rate reflects heat loss efficiency from the heated region, this parameter serves as an indicator of cellular heat retention capacity; faster relaxation corresponds to reduced heat retention. We then visualized the intracellular temperature changes under conditions that enhance endogenous heat production. The mitochondrial uncoupler FCCP disrupts the coupling between electron transport and ATP synthesis, thereby increasing electron transport activity and associated heat production in the mitochondria. As previously observed using independent methods^6,35^, FCCP treatment induced a temporal increase in the intracellular temperature in control cells (Fig. 3g and S5). In contrast, in cholesterol-depleted cells, the FCCP-induced increase in intracellular temperature was significantly suppressed, despite an unchanged mitochondrial membrane potential (Figs. S4a and 3g). These findings suggest that disturbing the plasma membrane decreases the ability of cells to retain metabolically generated heat, thereby increasing exothermic heat flow, as measured using DSC. In summary, we demonstrated that the plasma membrane contributes to cellular heat retention by limiting heat dissipation. Through these quantitative analyses, we discovered that not all metabolically generated heat is released from living cells. In other words, by resolving the cellular heat balance, we demonstrated that the cells actively limited the dissipation of heat across the cell boundary, thereby retaining a portion of the generated heat.

### Inhibition of transbilayer mobility of cell membrane lipids confers cells with heat retention ability

We investigated the mechanism by which the cell membrane suppresses heat dissipation. Cholesterol, which is removed from membranes by MβCD, is a key determinant of membrane identity^36^, as illustrated in Fig. 4a. Therefore, we examined the contribution of each mode to the heat dissipation rate from the cells.

**Figure 4.**
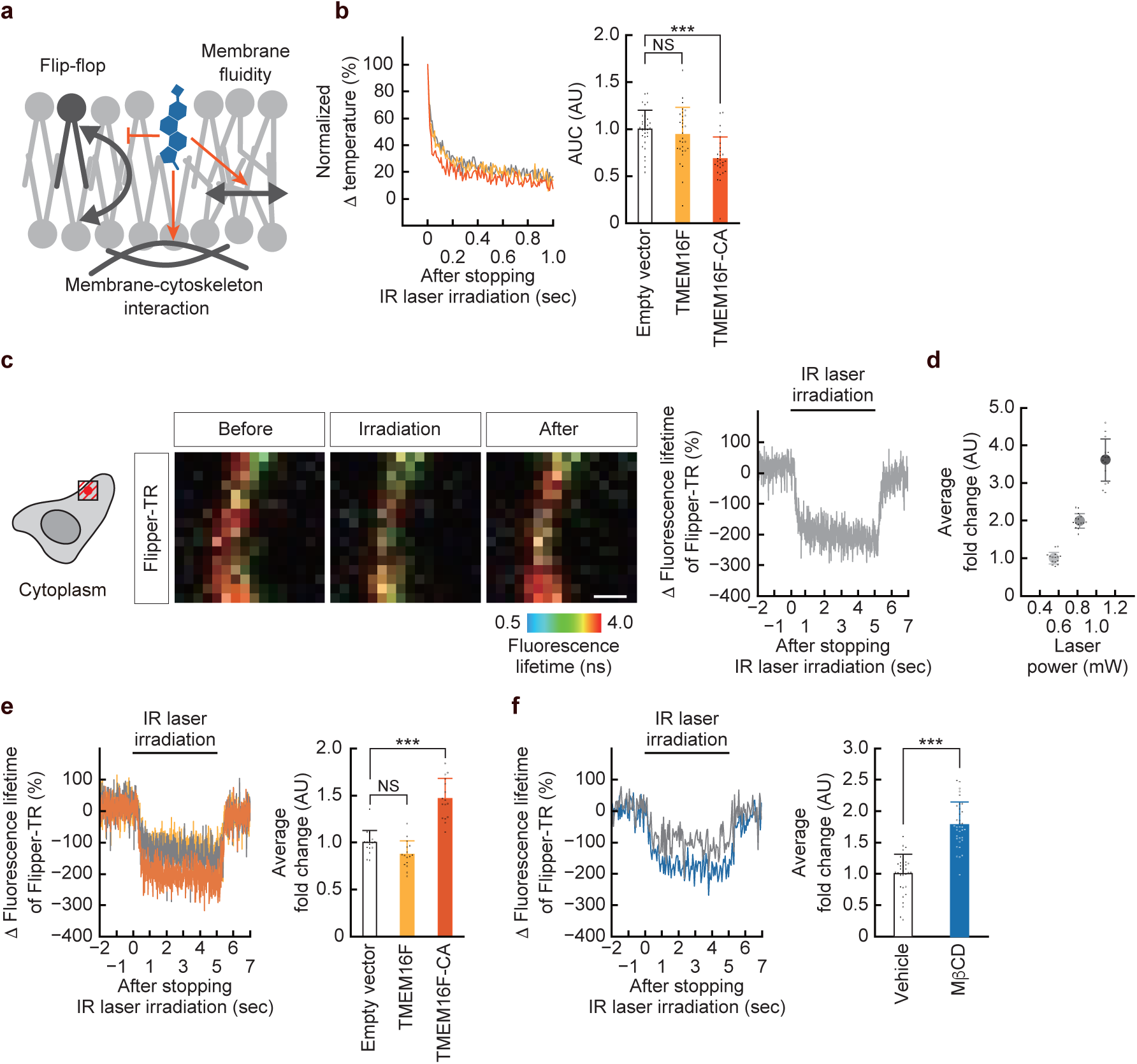
Identification of the molecular mechanism underlying cell membrane-dependent non-diffusive heat transfer inside cells. **a.** Three modes by which cholesterol contributes to plasma membrane integrity: reduction of membrane fluidity (lateral diffusion of biomolecules), reinforcement of membrane-cytoskeleton interactions, and suppression of spontaneous phospholipid flip-flop (transbilayer molecular movement). **b.** Temperature relaxation after stopping IR laser irradiation (0.8 mW for 5 s) in cells overexpressing the phospholipid scramblase TMEM16F. Traces were normalized to the temperature at the time point when irradiation was stopped. Data are shown as means (left). The relaxation rate, an index of intracellular heat transfer, was quantified as the area under the relaxation curve (AUC). Data are shown as means + s.d. (right) (*n* = 32 [Vector], 30 [TMEM16F], and 28 [TMEM16F-CA] cells). **c.** Tracking of the average Flipper-TR fluorescence lifetime within a square area measuring 10 μm on each side and centered on the heating point during heating of the juxtamembrane cytoplasmic region by IR laser irradiation (0.8 mW for 5 s). Changes in fluorescence lifetime (Δ) relative to the preheating baseline were evaluated. Data are shown as means (*n* = 15 cells). Scale bar: 2 μm. **d.** Relationship between IR laser power and Δ Fluorescence lifetime of Flipper-TR during localized heating for 5 s. Data are shown as means + s.d. (*n* = 15 cells). **e, f.** Δ Fluorescence lifetime of Flipper-TR at the plasma membrane during IR laser irradiation (0.8 mW for 5 s) in cells overexpressing TMEM16F **(e)** or in cells treated with MβCD **(f)**. Mean traces are shown (left in each panel). For bar graphs, the maximum Δ Fluorescence lifetime in each trace was extracted, and the values are shown as fold changes relative to the average value of cells transfected with the empty vector. Data are shown as means + s.d. (right in each panel) (e, n = 16 [Vector], 16 [TMEM16F], and 16 [TMEM16F-CA] cells; f, *n* = 33 [Vehicle] and 35 [MβCD] cells). Vehicle indicates treatment with PBS. CA, constitutively active mutant. \*\*\**P* < 0.001, NS indicates not significant.

First, we examined whether the cholesterol depletion–induced enhancement of cellular heat dissipation could be attributed to the disruption of the interaction between the plasma membrane and the underlying actin cortex. The actin-depolymerizing agent cytochalasin D (CytoD), which perturbs membrane integrity by disrupting the cortical actin network, did not affect the intracellular heat relaxation rate (Fig. S6a). Cholesterol also increases membrane fluidity, thereby enhancing the lateral diffusion of membrane-residing molecules. Membrane fluidity is influenced by the composition of the phospholipid acyl chains as well; phospholipids containing unsaturated fatty acids generally confer higher fluidity. Therefore, we used CAY10566^37^, an inhibitor of unsaturated fatty acid biosynthesis that reduces the abundance of unsaturated phospholipid acyl chains and consequently decreases membrane fluidity. However, CAY10566 treatment did not significantly alter the intracellular heat relaxation rate (Fig. S6b).

Next, we focused on another type of molecular motion within the membrane that is distinct from lateral diffusion: the transbilayer movement of lipids, known as flip-flop. Membrane cholesterol strengthens hydrophobic interactions among phospholipid acyl chains, suppressing spontaneous flip-flop^38^. Furthermore, because flip-flop requires high activation energy, it primarily occurs through bidirectional lipid movement mediated by phospholipid scramblases^39^. To mimic cholesterol-depleted membranes, in which spontaneous flip-flop increases due to disrupted membrane integrity, we overexpressed the scramblase TMEM16F in HeLa cells to artificially induce transbilayer lipid movement^40^. TMEM16F is normally inactive but becomes activated by elevations in cytosolic Ca^2+^, whereas the D409G mutant is constitutively active in a Ca^2+^-independent manner (TMEM16F-CA). Cells overexpressing wild-type TMEM16F exhibited intracellular heat transfer rates comparable to those of control cells. In contrast, TMEM16F-CA overexpression significantly accelerated the intracellular heat transfer, reducing the AUC by 31% (Fig. 4b). These findings suggest that suppressed transbilayer lipid movement is responsible for limiting heat dissipation from cells.

To clarify the causal relationship between flip-flop and heat dissipation from cells, we performed direct observations of the molecular dynamics at the plasma membrane during localized heating using an IR laser. Flipper-TR indicates the state of lipid packing within the membrane, and its fluorescence lifetime decreases in loosely packed membranes^41^. The fluorescence lifetime of Flipper-TR decreased in the plasma membrane upon localized heating of the juxtamembrane cytoplasmic region by IR laser irradiation. This demonstrates that lipid packing loosens during heat transfer from the inside to the outside of the cell (Fig. 4c). The magnitude of the change in membrane packing induced by IR laser irradiation was also confirmed to be quantitative, as evidenced by its dependence on the laser power (Fig. 4d). Loose lipid packing reflects weakened intermolecular interactions, leading to enhanced molecular motion, whereas tighter packing strengthens interactions and restricts molecular motion. Thus, we reasoned that this system would enable the quantitative assessment of membrane lipid mobility at the plasma membrane during intracellular heat transfer. In cells overexpressing wild-type TMEM16F, the decrease in the fluorescence lifetime of Flipper-TR upon localized heating was comparable to that in control cells. Conversely, overexpression of TMEM16F-CA enhanced the Flipper-TR response during IR laser irradiation (Fig. 4e). These results clearly demonstrate that flip-flop activation promotes membrane molecular motion during heat transfer from the inside to the outside of the cell. Furthermore, cholesterol-depleted cells exhibited increased membrane molecular motion, similar to that of TMEM16F-CA–expressing cells, during IR laser irradiation (Fig. 4f). This indicates that MβCD-induced membrane perturbation accelerates intracellular heat transfer by enhancing the transbilayer movement of membrane lipids. Based on our experiments, we conclude that heat dissipation from cells is governed by the conversion of thermal energy into the movement of membrane lipids, particularly the flip–flop of lipids across the transbilayer at the cell boundary. Cholesterol in the membrane helps restrain this process, which gives mammalian cells the ability to retain metabolically generated heat.

### Membrane dynamics create localized high-temperature regions within cells

Finally, we investigated the relationship between membrane dynamics, identified as key suppressors of cellular heat dissipation, and the formation of intracellular temperature distributions using FPT-based nanothermometry. As previously reported, the fluorescence lifetimes of FPT are heterogeneously distributed under steady-state conditions, indicating the presence of local high temperatures^6^. However, in cells overexpressing TMEM16F-CA, regions with long fluorescence lifetimes were not observed, as reflected by a marked reduction in lifetime variation (Fig. 5a). Similarly, the spatial pattern of FPT lifetimes in MβCD-treated cells exhibited notable homogeneity, with reduced lifetime variation (Fig. 5b). These findings indicate that, under both conditions, accelerated intracellular heat transfer prevents locally generated heat from being retained, thereby abolishing the formation of local high temperature regions. This figure therefore provides a direct, single-cell-level visualization of disrupted heat balance, fully consistent with our overall findings, including the calorimetric data.

**Figure 5.**
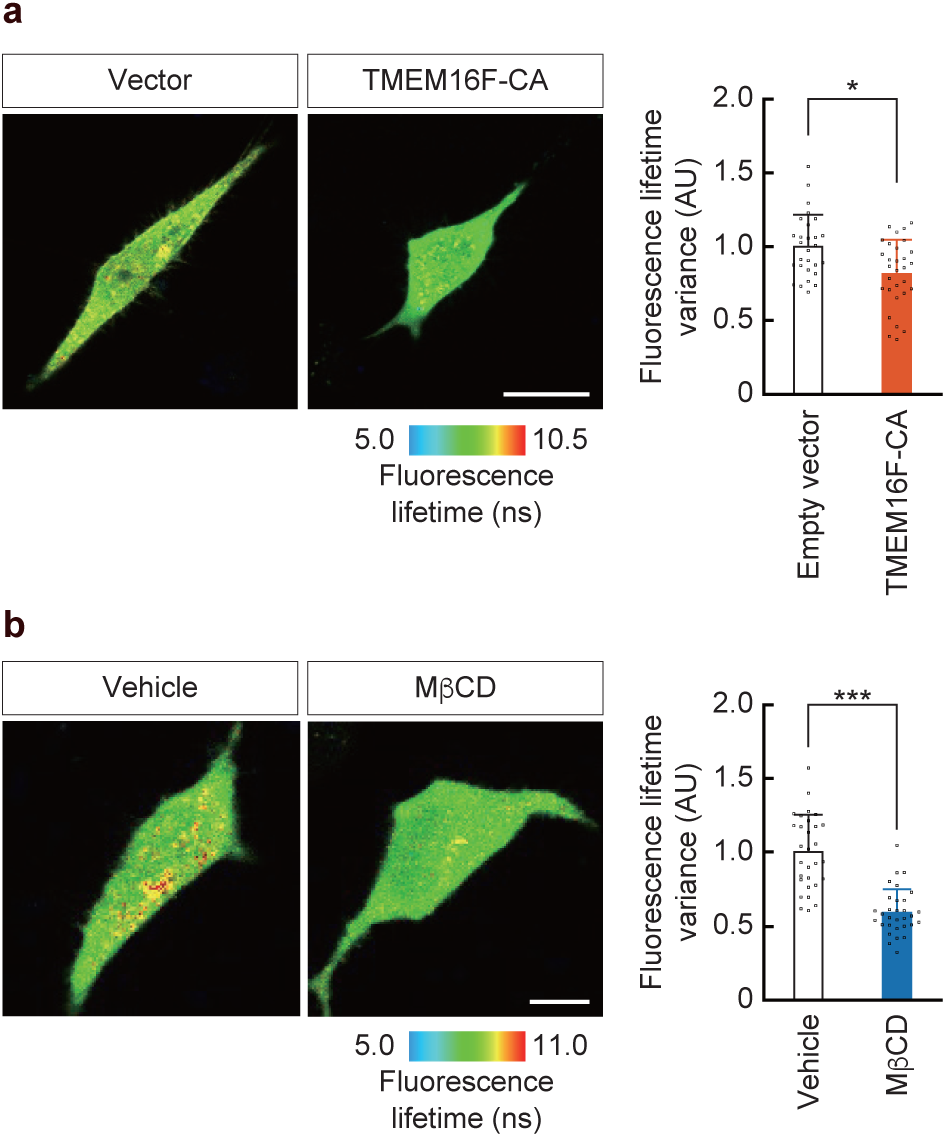
Temperature mapping in cells with disrupted membrane lipid dynamics. **a.** Temperature distribution in HeLa cells overexpressing a constitutively active mutant (CA) of the phospholipid scramblase TMEM16F. **b.** Temperature distribution in MβCD-treated cells. The heterogeneity of temperature distribution was quantified by the variance of FPT fluorescence lifetimes within cells (a, *n* = 29 [Vector] and 31 [TMEM16F-CA] cells; b, *n* = 30 [Vehicle] and 31 [MβCD] cells). Vehicle indicates treatment with PBS. Scale bars: 20 μm. \**P* < 0.05, \*\*\**P* < 0.001.

## Discussion

Conventionally, it has been assumed that because heat rapidly diffuses within cells, the flow of heat generated through catabolic reactions equals that of the heat dissipated and that all metabolically generated heat is immediately released from the inside to the outside of living cells. However, our previous study showed that intracellular heat transfer is slow and non-diffusive and that cells utilize intracellularly generated heat for cellular functions. Moreover, to the best of our knowledge, the direct cellular heat flux measurements underlying this assumption have not been experimentally verified. In this study, we employed DSC to successfully detect total exothermic heat flow derived from cellular catabolism within the physiological temperature range and found that this metabolic heat signature was distinctly different in character from the transient heat flow detected when biopolymers undergo state changes at specific (high) temperatures, such as during thermal denaturation (Fig. 1). We hypothesized that the cell membrane is responsible for regulating heat dissipation from the cells. Treatment with MβCD, a compound capable of effectively perturbing the cell membrane, resulted in a significant increase in heat output. This observation suggests that the cell membrane serves as the molecular basis for non-diffusive heat retention within cells, thereby preventing the dissipation of heat to the extracellular environment (Fig. 2). While the increase in heat dissipation could in principle arise from enhanced heat production activity or reduced heat retention capacity, analyses of organelle functions and single-cell thermometry demonstrated that MβCD specifically impaired the latter without affecting heat production activity. In addition, using our quantitative local heating technique, we directly observed that MβCD accelerates heat transfer within the cytosol, confirming that the elevated heat dissipation results from increased intracellular heat transfer and reduced heat retention capacity (Fig. 3). Furthermore, visualization of molecular dynamics at the plasma membrane revealed that heat dissipation is accompanied by enhanced molecular motion of membrane lipids, which was further enhanced by MβCD treatment. We identified the flip-flop as the motion responsible for the heat transfer process (Fig. 4). Finally, we demonstrated that cell membrane dynamics are essential for maintaining localized high-temperature regions within cells (Fig. 5). Collectively, these findings indicate that a portion of the non-diffusive heat energy remains within the cell, suggesting that cells actively retain metabolically generated heat and economically harness it for cellular functions. We propose that the molecular dynamics of the cell membrane act as an energetic barrier to heat dissipation from cells.

Similar to all the existing approaches, nanothermometry within cells, in which thousands of heat flux reactions occur with a highly heterogeneous spatial distribution, cannot capture or resolve the global heat balance in living cells. To overcome this limitation, we employed calorimetry, specifically DSC, in the present study. DSC enables the comprehensive quantification of cellular heat flow. Using our DSC system, we detected an exothermic heat flow of approximately 71 pW per HeLa cell at physiological temperature. This value is comparable to those previously reported using ITC on cells suspended in a culture medium^28,29^. However, earlier DSC-based studies reported smaller values of exothermic heat flow (e.g., 6.5 pW per cell)^24^. This discrepancy is likely due to differences in cell types and experimental conditions. In Loike’s study, the cells were suspended in a nutrient-poor buffer (5 mM glucose and dialyzed fetal bovine serum [FBS]). We also observed a reduction in exothermic flow under similar conditions (Fig. 1f). Our finding that living cells retain part of the metabolically generated heat (Fig. 2) demonstrates that calorimetrically measured heat output cannot be interpreted solely as a readout of heat-producing activity. Instead, the cellular capacity for heat retention must be considered. Using the intracellular heat transfer rates obtained from our quantitative single-cell temperature manipulation, we determined the contributions of heat production and dissipation from the signals measured by DSC. In other words, by integrating conventional calorimetry with state-of-the-art intracellular thermometry, we succeeded in resolving the cellular heat balance for the first time, which had previously remained inaccessible.

Discussions of subcellular temperature have long been based on the underlying assumptions that heat propagates within cells by conduction and that intracellular environments behave as thermodynamic equilibrium systems^16^. However, our previous study demonstrated that intracellular heat transfer does not rely solely on conduction^20^. Furthermore, experimental evidence of intracellularly retained heat, formed by energy conversion, provides direct proof that the intracellular environment is in a state of thermodynamic non-equilibrium. This provides new insights into how heat spreads within cells, including the non-diffusive heat transfer previously reported in our work. This unique intracellular thermodynamic environment creates several degrees of spatiotemporal temperature changes inside the cells, which our methods and various other nanothermometric approaches captured. Indeed, in cells where the membrane-dependent heat retention mechanism was impaired, such as after MβCD treatment or TMEM16F-CA overexpression, local high temperatures were no longer observed (Fig. 5). Nevertheless, the question of how the cell membrane influences the formation of intracellular local high temperatures remains unanswered, as MβCD primarily removes cholesterol from the plasma membrane, which is the outermost membrane of the cell^31^. Recent advances in electron and super-resolution microscopy have revealed that intracellular membranes are organized into networks through the cytoskeleton and membrane contact sites, forming the basis for intracellular material transport^42^. Therefore, it is conceivable that the physical state of the plasma membrane can modulate heat transport throughout the cell. However, further studies are required to elucidate this proposed mechanism.

Cells with enhanced flip-flop of membrane lipids exhibited an increased rate of intracellular heat transfer. This led us to conclude that the suppression of flip-flop is a key factor restricting heat dissipation from cells. In living cells, flip-flop is constitutively restrained by flippases, and scramblases remain inactive under normal conditions. However, during apoptosis, flip-flop is promoted through the proteolytic inactivation of flippases and caspase-dependent activation of scramblases. This results in the exposure of phosphatidylserine, a well-known “eat-me” signal^39^. We applied our quantitative local heating assay to examine whether the activation of flip-flop induced by pharmacologically triggered apoptosis affected cellular heat dissipation. Ionomycin, a Ca^2+^ ionophore that induces apoptosis by elevating cytosolic Ca^2+^^43^, increased the intracellular heat transfer rate (Fig. S7a). Similarly, staurosporine, a broad kinase inhibitor that triggers apoptosis by arresting the cell cycle^44^, accelerated intracellular heat dissipation in a time-dependent manner that paralleled apoptotic progression (Fig. S7b). These results confirm that activation of flip-flop promotes heat dissipation from the cells. Additionally, the capacity for heat retention inside the cells can be modulated during cellular processes. This raises the possibility that heat dissipation regulation may play a hitherto unrecognized physiological role.

In this study, we found that HeLa cells actively retained thermal energy. Importantly, we also confirmed that COS-7 cells treated with MβCD showed accelerated intracellular heat transfer (Fig. S8a) and enhanced exothermic heat flow (Fig. S8b), suggesting that membrane dynamics ubiquitously confer heat retention capacity to mammalian cells rather than in a cell-type–specific manner. We have previously identified thermal signaling, in which cells harness the heat they generate, such as glutamate-induced neural activity^14^ and NGF-induced neuronal differentiation^15^. These thermal signaling–dependent cellular phenomena are known to be suppressed by MβCD–induced plasma membrane perturbation^45,46^. Our results suggest that the reported effects of MβCD may stem from increased heat dissipation from cells and subsequent attenuation of thermal signaling. Given that thermal signaling can operate only when heat is effectively retained within the cell, these independent observations support our conclusion that a certain amount of heat energy is actively maintained inside the cells. Moreover, these results suggest that cellular heat retention may underlie a broad range of biological processes in which membrane dynamics play essential roles, such as cell migration, division, and fusion. Indeed, NIH-3T3 cells have been reported to increase their intracellular temperature during migration, implying that their capacity to retain heat may increase during this process^47^.

Because MβCD removes cholesterol from the plasma membrane, cholesterol appears to be an indispensable molecule for heat retention. However, some cell types contain far less membrane cholesterol than the HeLa cells used in this study, and some have been reported to have almost none^48^. Additionally, although we identified the suppression of membrane lipid flip-flop as the primary molecular mechanism that limits heat dissipation from living cells, a study on *Drosophila* cells showed that flip-flop occurs constantly at the plasma membrane^49^. These findings suggest two primary directions for future research. First, it is important to examine the differences in heat-retention capacity and thermal signaling contributions across cell types. For example, brown adipocytes, which consume lipids and generate heat, contain less cholesterol than white adipocytes, which store lipids^50^. This may reflect a physiological need for brown adipocytes to dissipate large amounts of generated heat to warm surrounding tissues in cold environments or a requirement to avoid excessive heat utilization (e.g., thermal signaling) within themselves. Second, it is important to identify factors beyond membrane dynamics that restrict heat dissipation. Inside cells, nucleic acids and proteins undergo structural transitions and complex assembly–disassembly cycles. The associated endothermic reactions either increase intracellular heat utilization or act as transient heat sinks. Consistent with this idea, we previously demonstrated that the heat transfer rate within the nucleus, where the rate is intrinsically slower than that in the cytoplasm, is accelerated upon treatment with the transcriptional inhibitor actinomycin D^20^. This suggests that structural transitions of nuclear nucleic acids influence cellular heat dissipation. Given the multitude of intracellular biopolymers capable of undergoing state changes, the establishment of high-throughput and refined screening systems will allow for the comprehensive identification of heat retention–related factors. These approaches will reveal novel molecular determinants that govern the fate of heat inside the cells.

Heat retention mediated by membrane lipid dynamics identified in this study represents an unrecognized cellular function of membrane lipids. In addition to our previous finding that membrane lipids regulate mitochondrial heat production^27^, we propose that they serve as master determinants of heat balance by controlling both the generation and retention of heat at the cellular level. Therefore, the physiological roles of membrane lipids should be reevaluated in light of their influence on intracellular temperature to deepen our understanding of their significance in biological systems. Notably, abnormalities in lipid metabolism are observed in a wide range of pathological conditions. However, in many cases, it remains unclear whether such changes represent primary causes or secondary consequences of the disease. The idea that membrane lipids regulate intracellular temperature and thermal signaling may provide a useful perspective for addressing this long-standing question. From a broader perspective, the composition of membrane lipids varies markedly across species, tissues, and even among cell types. We therefore hypothesize that this diversity in membrane lipids creates diversity in life activities by governing the cellular energy balance. This conceptual framework has the potential to trigger a paradigm shift not only in lipid biology but also across all life sciences.

## Methods

### Cell culture

HeLa (Riken, Wako, Japan) and COS-7 (Riken, Wako, Japan) cells were cultured in Dulbecco’s Modified Eagle’s Medium (DMEM) with 10% FBS supplemented with penicillin–streptomycin, L-glutamine, sodium pyruvate, and nonessential amino acids at 37°C in 5% CO_2_. For live-cell imaging, cells were cultured in 35-mm glass-bottomed dishes (AGC Techno Glass, Shizuoka, Japan), and the medium was replaced with a phenol red-free culture medium containing HEPES buffer (2 mL) before live-cell imaging. All solutions were purchased from Thermo Fisher Scientific (Waltham, MA, USA). The expression plasmids (pcDNA3.1-iRES-mCherry, TMEM16F-iRES-mCherry, and TMEM16F-CA-iRES-mCherry) were generously provided by Dr. Yuji Hara (University of Shizuoka). Cells were transfected with the plasmids using Lipofectamine (Thermo Fisher Scientific) for 3 and 24 h prior to experimentation. Rotenone, antimycin A, cytochalasin D, ionomycin, staurosporine, and dimethyl sulfoxide (DMSO) were purchased from WAKO (Osaka, Japan) and dissolved in DMSO. Carbonyl cyanide-p-trifluoromethoxyphenylhydrazone (FCCP) and CAY10566 were purchased from Sigma-Aldrich (Burlington, MA, USA) and dissolved in DMSO. 2-deoxy-D-glucose (2-DG, WAKO) and Methyl-β-cyclodextrin (MβCD, Sigma-Aldrich) were dissolved in PBS (PBS, WAKO).

### Preparation of the cell extract

HeLa cell pellets were collected and resuspended in a buffer (30 mM HEPES [pH 7.4], 75 mM sucrose, 225 mM mannitol, 0.1 mM ethylene glycol tetraacetic acid [EGTA]). After the addition of protease inhibitor cocktail (EDTA-free, Nacalai Tesque, Inc, Kyoto, Japan), the cell suspension was lysed using Dounce Tissue Grinder (Wheaton, Millville, NJ), followed by three rounds of centrifugation (600 ×*g*, 10 min, 4°C). Following further centrifugation (10,000 ×*g*, 10 min, 4°C), the crude mitochondrial fraction was removed from the supernatant. The protein concentration in the cell extracts was quantified using a BCA Protein Assay Kit (Thermo Fisher Scientific) after treatment with 1% Triton X-100 (Sigma-Aldrich).

### Measurement of glucose and lactate levels in the conditioned medium

Glucose and lactate concentrations in the conditioned medium were determined using the Glucose Assay Kit-WST and Lactate Assay Kit-WST (Dojindo, Kumamoto, Japan), respectively, according to the manufacturer’s instructions. Briefly, cells were seeded in 6-cm dishes in normal DMEM containing 25 mM glucose and cultured for 24 h. Then, the conditioned medium was collected and centrifuged at 1,000 ×*g* for 10 min. The supernatant was incubated with the reaction reagent, and absorbance was measured at 450 nm using a spectrophotometer (UH5200, HITACHI, Tokyo, Japan).

### Manipulation of intracellular cholesterol content

MβCD was dissolved in PBS to a concentration of 250 mM. Cells were incubated in FBS free-culture medium with 10 mM MβCD or PBS for 30 min at 37°C. MβCD-cholesterol inclusion complexes (MβCD + Chol) in Figs. 3b and 4f were generated by mixing a cholesterol suspension with an MβCD solution. In brief, cholesterol was added to MβCD solution at a concentration of 0.1 g mL^−1^. This saturated MβCD–cholesterol solution was incubated at 37°C overnight. Immediately before use, the solution was filtered through a 0.22-µm syringe filter (Millipore) to remove excess cholesterol crystals. This solution was added to the medium at a final MβCD concentration of 10 mM.

### Microcalorimetry

Microcalorimetry was based on DSC and performed using Nano DSC (TA Instruments). The device was equipped with three sample cells and a reference cell. The sample cells were filled with 700 µL of cell suspension, cell extract, or protein, while the reference cell contained 700 µL of the corresponding solvent. Initially, both the sample and reference cells were cooled to 5°C and equilibrated for 10 min. The 1^st^ scan was then conducted by heating from 5°C to 85°C at a rate of 0.5–2.0°C min^−1^. Following the 1^st^ scan, the temperature of these cells was cooled back to 5°C, equilibrated for 10 min, and subjected to a 2^nd^ scan at the same scan rate. Further scanning was performed (Fig. S1a, 3^rd^ scan). For the measurement of live cells, 1 × 10^7^ cells were used per run. For cell extracts, a scan was conducted using 5 mg of protein. In Fig. 1d, culture media with or without the same number of cells as in the living cell analysis were pre-heated at 85°C using a heat block and scanned in the sample cells and reference cell, respectively. In Fig. 3b, cells were trypsinized and incubated in 10 mM MβCD- or PBS containing-culture medium without FBS at 37°C for 30 min. Prior to DSC analysis, the cells were washed with PBS and resuspended in an FBS free-medium.

### Intracellular cholesterol imaging

Intracellular cholesterol was visualized using a Cholesterol Cell-based Detection Assay Kit (Cayman Chemical), according to the manufacturer’s instructions. Briefly, cells were fixed in Cell-Based Fixative Solution at 25°C for 10 min. After 5 min-wash with Cell-Based Assay Wash Buffer, cells were treated with Fillipin III solution at 25°C for 30 min and washed for 5 min twice with the wash buffer. Fluorescence images were acquired using a microscope (Axio-Observer Z1) with a 20 × objective lens. Fillipin III fluorescence was measured at 340 nm. The emissions were measured at 387/15 nm. The fluorescence intensity was quantified using ImageJ software. The average value of each cell was calculated.

### Mitochondrial membrane potential imaging

Mitochondrial membrane potential was evaluated using JC-1 (Dojindo Molecular Technologies), a lipophilic membrane-permeant cation that selectively enters the mitochondria and exists in a monomeric (green fluorescence) or aggregated form (red fluorescence) upon mitochondrial hyperpolarization. Cells were incubated with JC-1 (5 µM) for 30 min at 37°C and were washed twice with PBS. During microscopic observation, the cells were maintained in an FBS-free culture medium. Fluorescence images were obtained using a TCS SP8-FALCON confocal laser-scanning microscope (Leica Microsystems, Wetzeler, Germany) with a laser (552 and 488 nm) and an HC PL APO 63×/1.40 Oil CS2 objective (Leica Microsystems). Fluorescence was measured from 560 to 700 nm and from 500 to 700 nm, respectively. Ratiometric images (ex561/ex488) were analyzed using the ZEN software (Zeiss) and quantified using ImageJ. The average value of each cell was calculated.

### Intracellular Ca^2+^ imaging

Cells were seeded onto poly-L-lysine-coated coverslips and incubated overnight. The following day, cells were loaded with 5 µM Fura-2 AM (Dojindo Molecular Technologies) in culture medium at 37 °C for 40 min, followed by washing with HEPES-buffered saline (HBS) containing 2mM EGTA (HBS/EGTA). The coverslips were then transferred to a perfusion chamber mounted on an Axio Observer Z1 microscope (Zeiss, Oberkochen, Germany) at room temperature. Time-lapse imaging was performed at 10-s intervals. Cells were initially perfused with HBS/EGTA for 2 min, after which the solution was replaced with HBS/EGTA containing 3 µM Tg for 3 min. Subsequently, cells were exposed to HBS supplemented with 2 mM CaCl_2_ and 3 µM Ca²⁺ ionophore ionomycin. Ratiometric imaging (F340/F380) of Fura-2 fluorescence was conducted using the Physiology software (Zeiss). To account for cell-to-cell variability in the Fura-2 maximum response, fluorescence ratios were normalized to the ionomycin-induced maximum. Tg-induced cytosolic Ca²⁺ elevation was quantified as the difference between the peak Fura-2 ratio prior to ionomycin application and the ratio measured at 1 min after the start of imaging.

### Quantification of intracellular ATP content

The intracellular ATP content was measured using the ATP Assay Kit-Luminescence (Dojindo Molecular Technologies) according to the manufacturer’s instructions. Briefly, cells were wash with PBS and centrifuged at 3,000 ×*g* for 5 min at 4°C. The cell pallet was resuspended in PBS and incubated with Working solution in a 96-well white plate for 10 min at 25°C under dark conditions. Using Spark 10M (TECAN), the luminescence was measured at 5-s intervals. The protein concentration in the cell suspension was quantified using a BCA Protein Assay Kit (Thermo Fisher Scientific) after treatment with 1% Triton X-100.

### Fluorescence lifetime imaging

Cells were cultured in a 35-mm glass-bottomed dish in culture medium. For intracellular temperature imaging, cells were incubated in 150 µL of 5% glucose solution containing 0.02% cell-permeable FPT at 25°C for 10 min and washed twice with PBS. For imaging of biomolecular dynamics in cell membrane, cells were incubated in culture medium containing 3 µM Flipper-TR at 37°C for 15 min, followed by microscopic analysis without washout. Time-correlated single-photon counting-based FLIM was performed on a TCS SP8-FALCON confocal laser-scanning microscope (Leica Microsystems) with a pulsed diode laser (PDL 800-B, 470 nm; PicoQuant, Berlin, Germany) at a repetition rate of 20 MHz, and the emission from 500 to 640 nm was obtained through the described objective with 1–18.2 zoom factors and a binning procedure (factor: 1 or 2) in a 18×18–512×512-pixel format for 0.7–30.0 s. The temperature response curves for the temperature mapping with FPT in single cells were obtained by approximating the relationship between the averaged fast fluorescence lifetime (τ_FAST_) of FPT and the temperature to the third-degree polynomials (coefficients of determination for both *R*^2^ = 0.99):

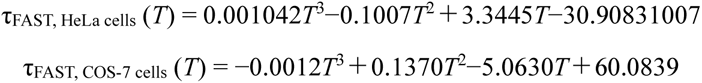

where *T* and τ_FAST_ (*T*) represent the temperature (°C) and the fast fluorescence lifetime (ns) at *T* °C, respectively. The temperature distribution was calculated from the standard deviation of τ_FAST_.

### Transient heating with IR laser irradiation

An IR laser (FRL-DC 1480 nm, maximum power, 3 W; IRE-Polus, MA, USA) was installed on a TCS SP8-FALCON confocal laser-scanning microscope (Leica Microsystems) using the IR-LEGO system (Sigma Koki, Tokyo, Japan) for the local heating of living cells. An HC PL APO 63 ’1.40 N.A. OIL CS2 objective (Leica Microsystems) was used to focus the IR laser and image the fluorescence lifetime of specimens; the diameter of the focused area in the IR laser irradiation was estimated to be 1.29 µm. An electronically controlled shutter (SSH-C4B; Sigma Koki) was used to start and stop IR laser irradiation. The irradiation duration was 5 s. The timing of stopping heating was determined analytically based on the rapid drop in FPT or Flipper-TR fluorescence intensity. Because shutter-controlled heating and image acquisition were not perfectly synchronized, the first frame after the fluorescence lifetime drop was excluded, as it may not have accurately reflected the temperature.

### Comparison of the time course of temperature relaxation

The time course of the temperature relaxation normalized by the temperature increase just before the cessation of heating by IR laser irradiation was analyzed. The AUC of normalized temperature relaxation for 0.1–1 s was also calculated, followed by statistical analysis. The points from 0–0.1 s were excluded from the calculation of AUC because the time required for the structural relaxation of FPT does not reflect the temperature.

## Acknowledgements

We thank Yuji Hara and Riku Nakanishi (University of Shizuoka) for kindly providing the TMEM16F expression vector and Masato Umeda (HOLO BIO Co., Ltd.) for critical comments on this manuscript. We are also grateful for the financial support from JST PRESTO and CREST (to K.O.), JST ACT-X (to A.M.), JSPS KAKENHI (18H03981, 20H05785, 24H02306, and 25K02236 to K.O.; 20J01234 and 22K20636 to A.M.), Life Science Foundation of Japan, and Brain Science Foundation.

## Author contributions

K.O. organized the study. A.M., K.O., and T.F. designed the study. A.M. performed all the imaging experiments and analyzed the data. A.M., T.S., and H.S. performed the DSC experiments and analyzed the data. K.O. and T.F. contributed to the interpretation of the results. A.M. and K.O. drafted the manuscript.

## Data availability

The data that support the findings of this study are available from the corresponding author upon reasonable request.

## Supplementary Note. Calculation of the total heat generated by cells through catabolic metabolism

Under the culture conditions used in this study, glucose is regarded as the sole substrate for catabolic metabolism. Therefore, we estimated the total amount of heat generated per cell through catabolism, based on the rate of glucose consumption in the culture medium and the associated reaction enthalpies.

During catabolism, glucose is oxidized to carbon dioxide (CO_2_) and water under aerobic conditions, yielding a theoretical maximum of 38 ATP molecules per glucose molecule (S1). Under anaerobic conditions, glucose is converted to lactate, which produces two ATP molecules per glucose molecule (S2). The heat generated in S1 was calculated from the heat released by complete oxidation of glucose and the free energy stored in the produced ATP as −2803 + 38 × 30.5 = −1644 kJ mol⁻¹, while the heat generated in S2 was calculated as −115 + 2 × 30.5 = −54 kJ mol⁻¹.

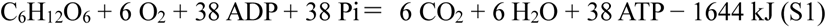

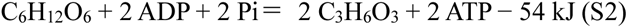

The relative contributions of aerobic and anaerobic metabolism were estimated from the amount of glucose consumed by cells from the culture medium, partitioned into total glucose consumption and the fraction metabolized anaerobically. The latter was calculated from the molar amount of lactate released into the medium, assuming that two lactate molecules were produced per molecule of glucose metabolized via anaerobic glycolysis. In this study, glucose consumption and lactate production were measured during a 24 h (1 d) culture period (Fig. 2). To calculate the glucose consumption rate per cell, *V* (mol day^−1^), we first considered the increase in cell number during the culture period (S3), where *n* denotes the cell number, *t* denotes time (days), and *m* is a cell-type-specific constant. From the initial condition at *t* = 0 with *n* = *n*_0_ and the value at *t* = 1 with *n* = *n*_1_, *n* and *m* can be described as follows (S4):

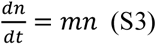

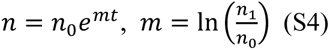

As the rate of change in glucose concentration in the culture medium, 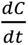, is given by *V* × *n*, the relationship is shown in S5. From the initial condition at *t* = 0, where *C* = *C*_0_, the value at *t* = 1 can be defined as shown in S6.

The aerobic glucose consumption rate (*v*_aero_) was obtained by substituting the molar amount of glucose metabolized aerobically, whereas substituting the molar amount of glucose metabolized anaerobically yields the anaerobic glucose consumption rate (*v*_anaero_). Finally, the total heat generated by the cells through glucose catabolism was estimated as *v*_aero_ × 1644 × 10^3^ + *v*_anaero_ × 54 × 10^3^ (J s^−1^).

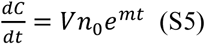

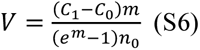

Notably, the efficiency of ATP synthesis during catabolic reactions is not 100%. Moreover, a fraction of the energy released by ATP hydrolysis may not be coupled to cellular functions and may instead be dissipated as heat. Conversely, as the thermodynamic state of ATP is influenced by the intracellular Mg²⁺ concentration and pH, the magnitude of the free energy stored in ATP may be greater inside living cells. Considering these factors, in practice, the amount of heat generated by the catabolic reactions described in S1 and S2 is expected to be higher or lower, respectively. Thus, the total heat generated in our calculations is regarded as an approximate estimate. In addition, the energy produced by catabolism can contribute to anabolic processes only through conversion to ATP. Accordingly, our analysis excludes the possibility that catabolically generated heat was directly coupled with anabolic metabolism.

---

**Supplementary Figure 1. Baseline trend in raw DSC curve.**

**a.** Raw DSC traces obtained from three repeated scans of the same HeLa cell sample.
**b.** Corrected DSC traces generated by subtracting the second scan from the first and the third scan from the second using the dataset shown in **a**.

**Supplementary Figure 2. Detection of heat flow associated with thermal denaturation of bovine serum albumin (BSA) using our DSC system.**

BSA dissolved in phosphate-buffered saline (PBS) at a concentration of 30 mg mL^−1^ was analyzed by DSC at scan rates of 0.5, 1.0, and 2.0 °C min^−1^ (*n* = 3).

In the DSC thermograms, mean traces are shown. Negative values on the y-axis represent exothermic heat flow, whereas positive values indicate endothermic heat flow. For bar graphs, the maximum endothermic signal in each DSC thermogram was extracted and defined as Max endo-flow. Data are shown as means + standard deviation (s.d.). The dotted lines indicate linear fitting. R indicates the correlation coefficient; p values were used to assess statistical significance, with *P* < 0.05 considered significant.

**Supplementary Figure 3. Quantification of cellular heat balance.**

HeLa cells were cultured for 24 h.

**a.** Change in HeLa cell number.
**b.** Glucose consumption in the medium.
**c.** Lactate release into the medium.
**d.** Comparison of heat generated in cells, calculated from catabolic metabolism (details shown in the Supplementary Note), with heat dissipated from cells (exothermic heat flow) at 37 °C and the maximum exothermic heat flow measured by DSC. Data are shown as means + standard deviation (s.d.) (*n* = 3).

\*\*\**P* < 0.001.

**Supplementary Figure 4. Effects of cholesterol depletion by MβCD on the functions of thermogenic organelles.**

**a.** Mitochondrial membrane potential in HeLa cells assessed using JC-1. Carbonyl cyanide *p*-trifluoromethoxyphenylhydrazone (FCCP), an uncoupler, was used as a positive control (5 µM for 15 min) for detecting changes in membrane potential. Data are shown as means + standard deviation (s.d.) (*n* = 75 [Vehicle], 84 [MβCD], and 81 [FCCP] cells). Scale bar: 50 µm.
**b.** Intracellular oxygen consumption rate (OCR). Oligomycin (Oligo), rotenone (Rot), and antimycin A (AA) were used as inhibitors of ATP synthase and mitochondrial respiratory chain complexes I and III, respectively. Data are shown as means + s.d. (*n* = 3).
**c.** Activity of sarco/endoplasmic reticulum Ca²⁺-ATPase (SERCA) assessed using Fura-2. Changes in the Fura-2 ratio from baseline were monitored as an indicator of cytoplasmic Ca²⁺ concentration. Traces were normalized to baseline and to the maximal response induced by ionomycin. Ionomycin treatment served as a positive control for the detection. Data are shown as means (left). The cytoplasmic [Ca²⁺] response to thapsigargin (Tg) was used as an index of SERCA activity. Data are shown as means + s.d. (right) (*n* = 88 [Vehicle], n = 84 [MβCD] cells).
**d.** Intracellular ATP levels. Treatment with rotenone and antimycin A (Rot/AA) was used as a positive control for detecting changes in ATP levels. Data are shown as means + s.d. (*n* = 3).

Vehicle indicates treatment with PBS. \*\**P* < 0.01, \*\*\**P* < 0.001, NS indicates not significant.

**Supplementary Figure 5. Tracking of intracellular temperature in HeLa cells during dimethyl sulfoxide (DMSO) treatment.**

The average intracellular temperature of whole cells was tracked after DMSO addition using FPT. ΔTemperature represents the change in temperature relative to the value before DMSO addition. Data are shown as means + standard deviation (s.d.) (*n* = 20 cells). NS indicates not significant.

**Supplementary Figure 6. Intracellular heat transfer in cells with disrupted membrane dynamics-related factors.**

**a, b.** Cytoplasmic temperature relaxation after stopping IR laser irradiation (0.8 mW for 5 s) in cells treated with cytochalasin D (CytoD; 400 nM for 3 h), an actin-depolymerizing agent **(a**), or CAY10566, an inhibitor of unsaturated fatty acid biosynthesis (1 µM for 16 h) **(b)**. Traces were normalized to the temperature at the time point when irradiation was stopped. Data are shown as means (left in each panel). The relaxation rate, an index of intracellular heat transfer, was quantified as the area under the relaxation curve (AUC). Vehicle indicates treatment with PBS. Data are compared with those obtained from living HeLa cells (Fig. 4f). Data are shown as means + s.d. (right in each panel) (a, *n* = 29 [Vector] and 25 [CytoD] cells; b, *n* = 29 [Vehicle] and 31 [CAY10566] cells). NS indicates not significant.

**Supplementary Figure 7. Intracellular heat transfer in cells undergoing apoptosis.**

**a, b.** Cytoplasmic temperature relaxation after stopping IR laser irradiation (0.8 mW for 5 s) in cells treated with ionomycin, a Ca²⁺ ionophore (0.5 µM for 5-25 min) **(a**), or staurosporine (STS), a broad kinase inhibitor (10 µM for the indicated durations) **(b)**. Traces were normalized to the temperature at the time point when irradiation was stopped. Data are shown as means (left in each panel). The relaxation rate was quantified as the area under the relaxation curve (AUC). Vehicle indicates treatment with DMSO. Data are shown as means + s.d. (right in each panel) (a, *n* = 23 [Vehicle] and 23 [Iono] cells; b, *n* = 28 [0 h], 27 [2 h], and 29 [4 h] cells). \*\**P* < 0.01, \*\*\**P* < 0.001, NS indicates not significant.

**Supplementary Figure 8. Effect of MβCD on heat retention-related phenotypes in COS-7 cells.**

**a.** Cytoplasmic temperature relaxation after stopping IR laser irradiation (0.8 mW for 5 s). Traces were normalized to the temperature at the time point when irradiation was stopped. Data are shown as means (left). The relaxation rate, an index of intracellular heat transfer, was quantified as the area under the relaxation curve (AUC). Data are shown as means + s.d. (right) (n = 28 [Vehicle] and 31 [MβCD] cells).
**b.** Exothermic heat flow measured by DSC (*n* = 3). In the DSC thermograms, mean traces are shown. Negative values on the y-axis represent exothermic heat flow, whereas positive values indicate endothermic heat flow. For bar graphs, the maximum exothermic signal in each DSC thermogram was extracted and defined as Max exo-flow. Data are shown as means + standard deviation (s.d.).

Vehicle indicates treatment with PBS. \*\*\**P* < 0.001.

