## Supplementary information for "Membrane Dynamics-Mediated Heat Retention Dictates Economical Intracellular Energy Flow"

$$\frac{dn}{dt} = mn \quad (S3)$$

$$n = n_0 e^{mt},\ m = \ln\left(\frac{n_1}{n_0}\right) \quad (S4)$$

As the rate of change in glucose concentration in the culture medium, $\frac{dC}{dt}$, is given by $V \times n$, the relationship is shown in S5. From the initial condition at $t = 0$, where $C = C_0$, the value at $t = 1$ can be defined as shown in S6.

The aerobic glucose consumption rate ($v_{aero}$) was obtained by substituting the molar amount of glucose metabolized aerobically, whereas substituting the molar amount of glucose metabolized anaerobically yields the anaerobic glucose consumption rate ($v_{anaero}$). Finally, the total heat generated by the cells through glucose catabolism was estimated as $v_{aero} \times 1644 \times 10^3 + v_{anaero} \times 54 \times 10^3$ (J s$^{-1}$).

$$\frac{dC}{dt} = Vn_0e^{mt} \quad \text{(S5)}$$

$$V = \frac{(C_1 - C_0)m}{(e^m - 1)n_0} \quad \text{(S6)}$$

Notably, the efficiency of ATP synthesis during catabolic reactions is not 100%. Moreover, a fraction of the energy released by ATP hydrolysis may not be coupled to cellular functions and may instead be dissipated as heat. Conversely, as the thermodynamic state of ATP is influenced by the intracellular $Mg^{2+}$ concentration and pH, the magnitude of the free energy stored in ATP may be greater inside living cells. Considering these factors, in practice, the amount of heat generated by the catabolic reactions described in S1 and S2 is expected to be higher or lower, respectively. Thus, the total heat generated in our calculations is regarded as an approximate estimate. In addition, the energy produced by catabolism can contribute to anabolic processes only through conversion to ATP. Accordingly, our analysis excludes the possibility that catabolically generated heat was directly coupled with anabolic metabolism.

Murakami et al. Supplementary figure. 1

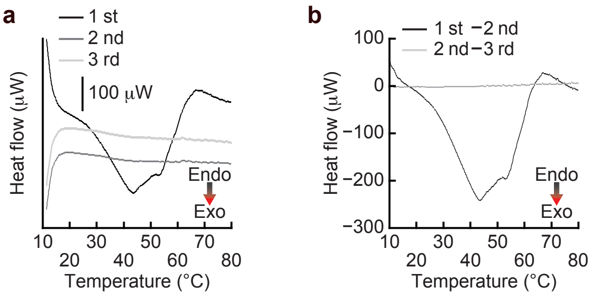

**Supplementary Figure 1. Baseline trend in raw DSC curve.**

**a.** Raw DSC traces obtained from three repeated scans of the same HeLa cell sample.

**b.** Corrected DSC traces generated by subtracting the second scan from the first and the third scan from the second using the dataset shown in **a**.

Murakami et al. Supplementary figure. 2

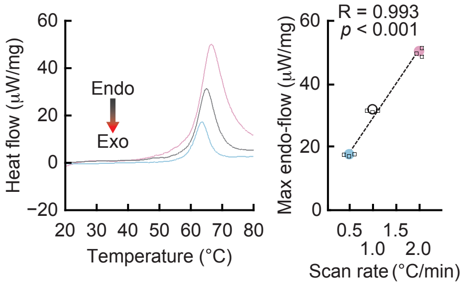

**Supplementary Figure 2. Detection of heat flow associated with thermal denaturation of bovine serum albumin (BSA) using our DSC system.**

BSA dissolved in phosphate-buffered saline (PBS) at a concentration of 30 mg $mL^{-1}$ was analyzed by DSC at scan rates of 0.5, 1.0, and 2.0 °C $min^{-1}$ ($n = 3$).

In the DSC thermograms, mean traces are shown. Negative values on the y-axis represent exothermic heat flow, whereas positive values indicate endothermic heat flow. For bar graphs, the maximum endothermic signal in each DSC thermogram was extracted and defined as Max endo-flow. Data are shown as means + standard deviation (s.d.). The dotted lines indicate linear fitting. R indicates the correlation coefficient; p values were used to assess statistical significance, with $P < 0.05$ considered significant.

Murakami et al. Supplementary figure. 3

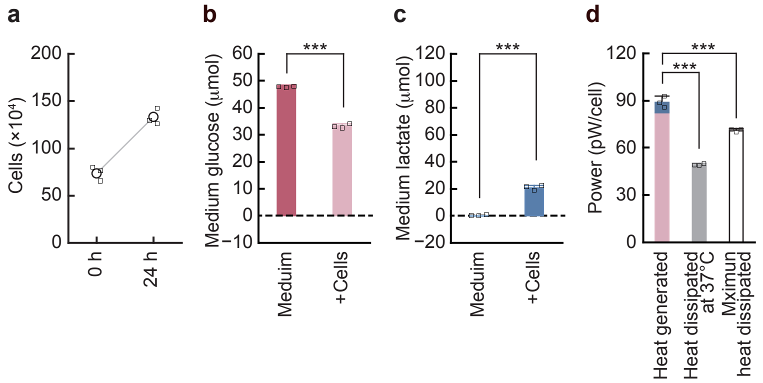

**Supplementary Figure 3. Quantification of cellular heat balance.**

HeLa cells were cultured for 24 h.

**a.** Change in HeLa cell number.

**b.** Glucose consumption in the medium.

**c.** Lactate release into the medium.

**d.** Comparison of heat generated in cells, calculated from catabolic metabolism (details shown in the Supplementary Note), with heat dissipated from cells (exothermic heat flow) at 37 °C and the maximum exothermic heat flow measured by DSC. Data are shown as means + standard deviation (s.d.) ($n = 3$).

***$P < 0.001$.

Murakami et al. Supplementary figure. 4

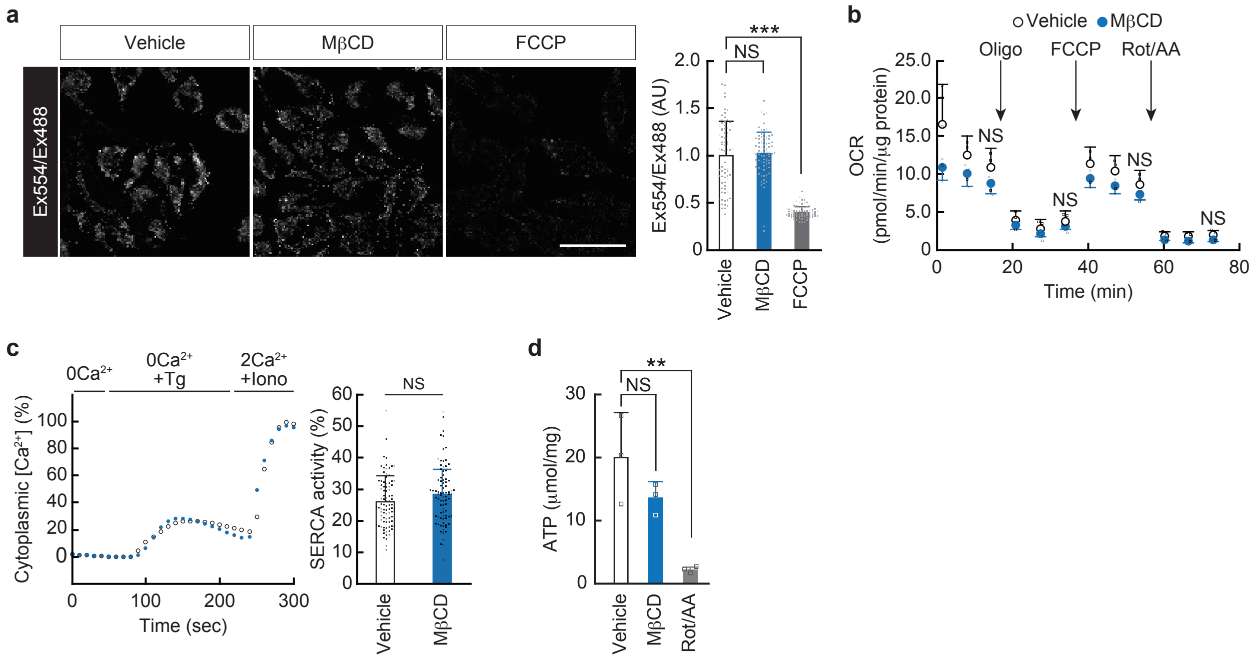

**Supplementary Figure 4. Effects of cholesterol depletion by MβCD on the functions of thermogenic organelles.**

**a.** Mitochondrial membrane potential in HeLa cells assessed using JC-1. Carbonyl cyanide *p*-trifluoromethoxyphenylhydrazone (FCCP), an uncoupler, was used as a positive control (5 µM for 15 min) for detecting changes in membrane potential. Data are shown as means + standard deviation (s.d.) ($n$ = 75 [Vehicle], 84 [MβCD], and 81 [FCCP] cells). Scale bar: 50 µm.

**b.** Intracellular oxygen consumption rate (OCR). Oligomycin (Oligo), rotenone (Rot), and antimycin A (AA) were used as inhibitors of ATP synthase and mitochondrial respiratory chain complexes I and III, respectively. Data are shown as means + s.d. ($n$ = 3).

**c.** Activity of sarco/endoplasmic reticulum $Ca^{2+}$-ATPase (SERCA) assessed using Fura-2. Changes in the Fura-2 ratio from baseline were monitored as an indicator of cytoplasmic $Ca^{2+}$ concentration. Traces were normalized to baseline and to the maximal response induced by ionomycin. Ionomycin treatment served as a positive control for the detection. Data are shown as means (left). The cytoplasmic [$Ca^{2+}$] response to thapsigargin (Tg) was used as an index of SERCA activity. Data are shown as means + s.d. (right) ($n$ = 88 [Vehicle], n = 84 [MβCD] cells).

**d.** Intracellular ATP levels. Treatment with rotenone and antimycin A (Rot/AA) was used as a positive control for detecting changes in ATP levels. Data are shown as means + s.d. ($n$ = 3).

Vehicle indicates treatment with PBS. **$P < 0.01$, ***$P < 0.001$, NS indicates not significant.

Murakami et al. Supplementary figure. 5

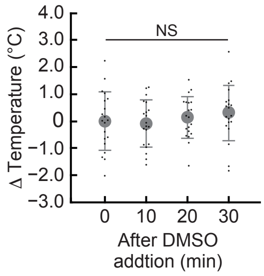

**Supplementary Figure 5. Tracking of intracellular temperature in HeLa cells during dimethyl sulfoxide (DMSO) treatment.**

The average intracellular temperature of whole cells was tracked after DMSO addition using FPT. ΔTemperature represents the change in temperature relative to the value before DMSO addition. Data are shown as means + standard deviation (s.d.) ($n$ = 20 cells). NS indicates not significant.

Murakami et al. Supplementary figure. 6

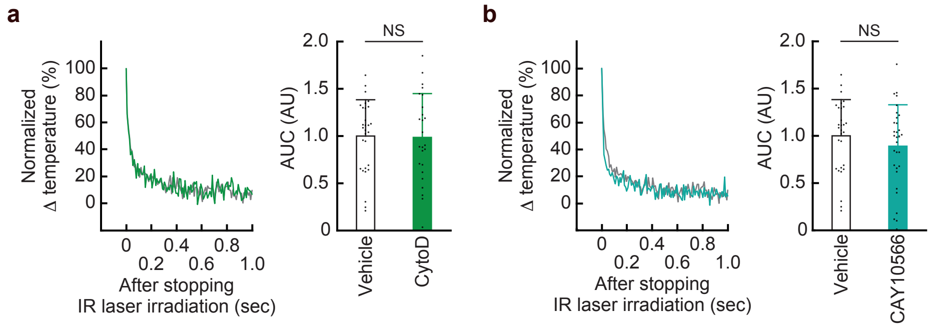

**Supplementary Figure 6. Intracellular heat transfer in cells with disrupted membrane dynamics-related factors.**

**a, b.** Cytoplasmic temperature relaxation after stopping IR laser irradiation (0.8 mW for 5 s) in cells treated with cytochalasin D (CytoD; 400 nM for 3 h), an actin-depolymerizing agent **(a)**, or CAY10566, an inhibitor of unsaturated fatty acid biosynthesis (1 μM for 16 h) **(b)**. Traces were normalized to the temperature at the time point when irradiation was stopped. Data are shown as means (left in each panel). The relaxation rate, an index of intracellular heat transfer, was quantified as the area under the relaxation curve (AUC). Vehicle indicates treatment with PBS. Data are compared with those obtained from living HeLa cells (Fig. 4f). Data are shown as means + s.d. (right in each panel) (a, $n$ = 29 [Vector] and 25 [CytoD] cells; b, $n$ = 29 [Vehicle] and 31 [CAY10566] cells). NS indicates not significant.

Murakami et al. Supplementary figure. 7

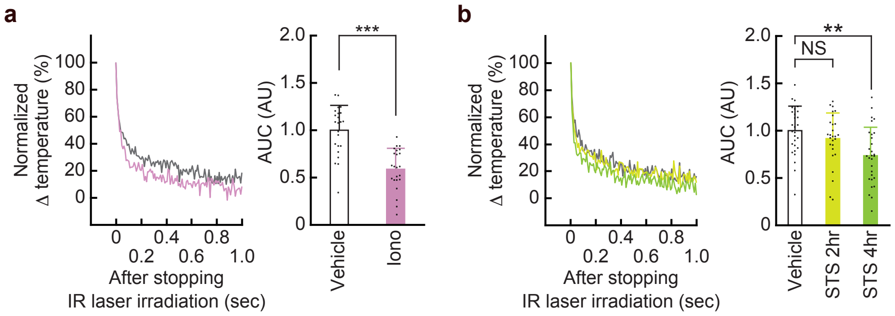

**Supplementary Figure 7. Intracellular heat transfer in cells undergoing apoptosis.**

**a, b.** Cytoplasmic temperature relaxation after stopping IR laser irradiation (0.8 mW for 5 s) in cells treated with ionomycin, a $Ca^{2+}$ ionophore (0.5 μM for 5-25 min) **(a)**, or staurosporine (STS), a broad kinase inhibitor (10 μM for the indicated durations) **(b)**. Traces were normalized to the temperature at the time point when irradiation was stopped. Data are shown as means (left in each panel). The relaxation rate was quantified as the area under the relaxation curve (AUC). Vehicle indicates treatment with DMSO. Data are shown as means + s.d. (right in each panel) (a, $n$ = 23 [Vehicle] and 23 [Iono] cells; b, $n$ = 28 [0 h], 27 [2 h], and 29 [4 h] cells). $^{**}P < 0.01$, $^{***}P < 0.001$, NS indicates not significant.

Murakami et al. Supplementary figure. 8

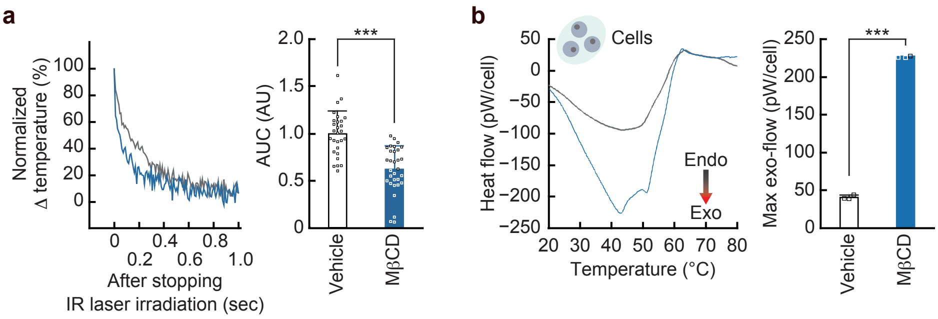

**Supplementary Figure 8. Effect of MβCD on heat retention-related phenotypes in COS-7 cells.**

**a.** Cytoplasmic temperature relaxation after stopping IR laser irradiation (0.8 mW for 5 s). Traces were normalized to the temperature at the time point when irradiation was stopped. Data are shown as means (left). The relaxation rate, an index of intracellular heat transfer, was quantified as the area under the relaxation curve (AUC). Data are shown as means + s.d. (right) (n = 28 [Vehicle] and 31 [MβCD] cells).

**b.** Exothermic heat flow measured by DSC ($n = 3$). In the DSC thermograms, mean traces are shown. Negative values on the y-axis represent exothermic heat flow, whereas positive values indicate endothermic heat flow. For bar graphs, the maximum exothermic signal in each DSC thermogram was extracted and defined as Max exo-flow. Data are shown as means + standard deviation (s.d.).

Vehicle indicates treatment with PBS. ***$P < 0.001$.
